# Genetic Diversity and Geographic Structure of Nordic Rye Landraces and Historic Varieties: Expected Patterns and New Insights

**DOI:** 10.64898/2026.11.25.769110

**Authors:** Teija T. Tenhola-Roininen, Pirjo K. Tanhuanpää, Maarit I. Heinonen, Hanna Haikka, Lidija Bitz

## Abstract

Rye has been an important food crop and source of nutrition in Finland for centuries and remains so today. Rye landraces and old varieties are still grown on farmers’ fields. Interest in local landraces and conservation varieties has grown during the last decade due to climate change and its negative effects on biodiversity and resilience. We explored genetic diversity and population structure in Nordic rye landraces, historic population varieties and reference materials using complementary SSR and SNP datasets. For comparative analysis of genetic diversity, the Finnish varieties ‘Riihi’ and ‘Reetta’, the German inbred line Lo7, a German hybrid variety, the Estonian variety ‘Elvi’, and the landrace ‘Sangaste’ were included as reference materials. Both SSR and SNP analyses revealed geographically associated genetic differentiation within an otherwise largely shared Nordic rye gene pool. Notably, the independently sampled Mustialan and NordGen accessions of the Norwegian ‘Tvengsberg’ landrace were each clearly differentiated from the remaining rye material in the SSR and SNP datasets, respectively. Finnish landraces (‘Aunus’, ‘Hermanni’, ‘Iivo’, ‘Joppe’, ‘Pääkkölä’, and ‘Taavetti’) showed broadly similar genetic profiles, although substantial admixture was evident. In the SSR analysis, the Estonian accessions ‘Sangaste’ and ‘Elvi’ clustered together with ‘Reetta’, ‘Riihi’, and Finnish landrace ‘Polvijärvi’, suggesting a closer genetic relationship among these accessions, although ‘Polvijärvi’ has previously been considered as a distinct Finnish landrace. Finnish variety ‘Hankkijan Jussi’ formed a separate cluster; this variety originates from a selection derived from the 60Co-irradiated Russian variety ‘Vjatka’. The modern inbred line Lo7 and an unnamed hybrid variety were included as outgroup controls and showed clear genetic divergence from the Fennoscandian landrace material. Although Nordic rye landraces are often assumed to share a common ancestry and may have been influenced by historical seed exchange among farming communities, both SSR and SNP analyses revealed geographically associated genetic differentiation. These results indicate that regional genetic differentiation has been maintained despite the outcrossing nature of rye and the potential for historical gene flow. The concordance between SSR and SNP analyses indicates that geographic differentiation remains detectable within a broadly shared Nordic rye gene pool

## Introduction

Cultivated rye (Secale cereale subsp. cereale) is an outcrossing diploid cereal species belonging to the family Poaceae and the tribe Triticeae (Evans 1976). Due to its allogamous breeding system, rye populations typically exhibit high levels of genetic diversity within populations. In Finland, winter rye has traditionally been the dominant type, although spring rye varieties also exist. The Nordic Genetic Resource Center (NordGen) currently maintains 146 Finnish rye accessions, including landraces, varieties, and breeding materials (GenBis database).

Rye has played an important role in Finnish agriculture and food culture for centuries (Ahokas 2009; Ahokas 2012). It became one of the most important cereal crops in Fennoscandia and remained widely cultivated because of its winter hardiness and reliable yield under northern conditions (Ahokas 2009; Ahokas 2012; Larsson et al. 2019). Finnish rye cultivation was historically associated with slash-and-burn agriculture, which contributed to the development of distinctive local landraces. Finnish settlers introduced these rye populations to central Sweden during the seventeenth century, and cultivation later spread to eastern Norway. Consequently, related rye landraces have been maintained across the Fennoscandian region for several centuries, providing a unique resource for studying long-term cultivation and regional adaptation.

In the 1970s, Norwegian ethnologist Per Martin Tvengsberg succeeded in germinating seven out of ten rye grains that had been found beneath the floorboards of an old drying barn (riihi) in Finnskogen, a forest region on the Norwegian-Swedish border. The resulting rye landrace was later named ‘Tvengsberg’ (Tvengsberg svedjerug) in recognition of his efforts to preserve and reintroduce the landrace (Natural History Museum and Botanical Garden 2026). Since 2017, the landrace has been cultivated on a very small scale for seed propagation following a slash-and-burn cultivation experiment carried out in Finland in 2004–2005 (Annika Michelson, pers. comm., 2023). This ‘Tvengsberg’ landrace is maintained in the NordGen genebank (GenBis database).

Today, rye landraces are cultivated only on a very small scale, mainly for conservation and cultural purposes. Nevertheless, they represent valuable genetic resources because they may harbor adaptive traits and genetic variation that have been lost in modern breeding materials. Genetic diversity in rye has been studied using a range of molecular markers. Among these, simple sequence repeats (SSRs) and single nucleotide polymorphisms (SNPs) are widely used. SSR markers are highly polymorphic and multi-allelic, making them particularly useful for assessing diversity within heterogeneous populations, whereas SNP markers enable high-throughput, genome-wide analyses (Hackauf and Wehling 2002; Targońska et al. 2016; Hagenblad et al. 2016; Schreiber et al. 2018). Recent studies have revealed extensive genetic diversity in rye landraces and traditional varieties, with most genetic variation occurring within, rather than among, populations.

The aim of this study was to assess the genetic diversity, population structure, and genetic relationships among Finnish rye landraces and conservation varieties maintained in cultivation and ex situ collections. Both SSR and SNP markers were used to characterize genetic variation and to evaluate the relative informativeness of these marker systems in heterogeneous rye populations.

## Materials and Methods

### Rye material

In total, sixteen rye conservation varieties and modern varieties/lines were selected for the study of the genetic diversity in open-pollinated rye populations (Table 1). The selection of rye conservation varieties was made in 2018. Seeds of eight out of 13 conservation varieties in Finland were obtained from farmers maintaining or cultivating them. In addition to these, five traditionally cultivated, two Estonian and three of Finnish origin, varieties were selected for the study. Selected landraces and traditional varieties have been cultivated in Finland for decades, except ‘Mustialan Tvengsberg’, which has been grown only since 2017.

**Table 1.**
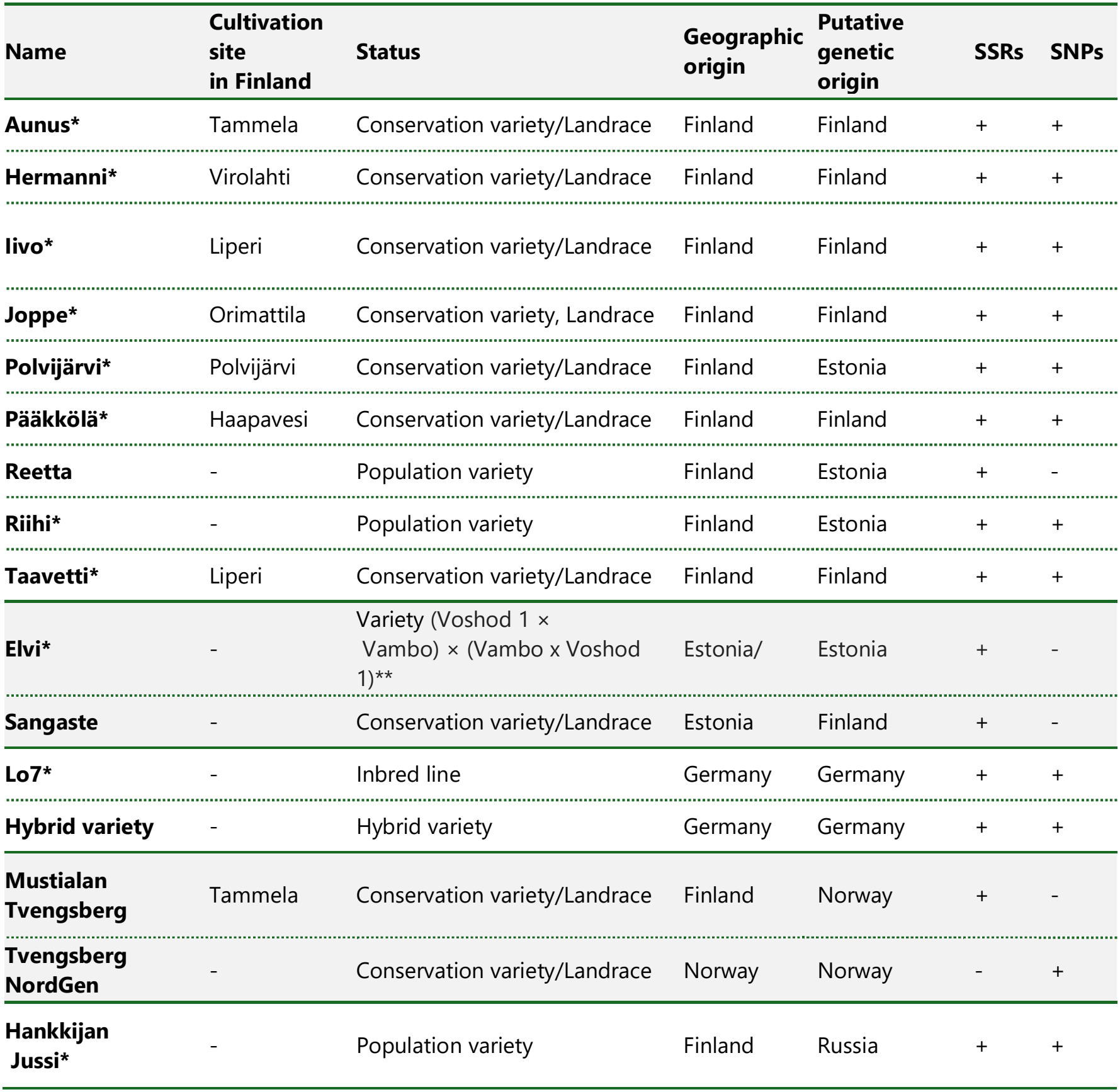
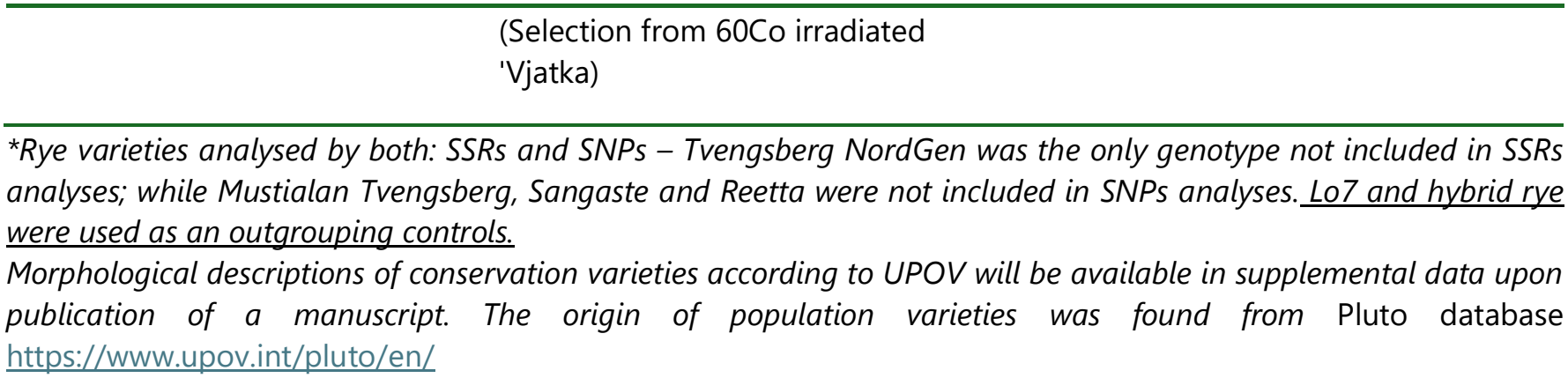
List of 16 rye varieties used in the study, including cultivation site (Finland, when available), status, origin, putative genetic origin, and SSR/SNP analyses. Geographic origin refers to the country or region where the accession was cultivated or collected, whereas putative genetic origin refers to the presumed origin of the germplasm from which the cultivar or population was developed, based on available oral, historical and pedigree information.

| Name | Cultivation site in Finland | Status | Geographic origin | Putative genetic origin | SSRs | SNPs |
|---|---|---|---|---|---|---|
| **Aunus\*** | Tammela | Conservation variety/Landrace | Finland | Finland | + | + |
| **Hermanni\*** | Virolahti | Conservation variety/Landrace | Finland | Finland | + | + |
| **Iivo\*** | Liperi | Conservation variety/Landrace | Finland | Finland | + | + |
| **Joppe\*** | Orimattila | Conservation variety, Landrace | Finland | Finland | + | + |
| **Polvijärvi\*** | Polvijärvi | Conservation variety/Landrace | Finland | Estonia | + | + |
| **Pääkkölä\*** | Haapavesi | Conservation variety/Landrace | Finland | Finland | + | + |
| **Reetta** | - | Population variety | Finland | Estonia | + | - |
| **Riihi\*** | - | Population variety | Finland | Estonia | + | + |
| **Taavetti\*** | Liperi | Conservation variety/Landrace | Finland | Finland | + | + |
| **Elvi\*** | - | Variety (Voshod 1 × Vambo) × (Vambo x Voshod 1)** | Estonia/ | Estonia | + | - |
| **Sangaste** | - | Conservation variety/Landrace | Estonia | Finland | + | - |
| **Lo7\*** | - | Inbred line | Germany | Germany | + | + |
| **Hybrid variety** | - | Hybrid variety | Germany | Germany | + | + |
| **Mustialan Tvengsberg** | Tammela | Conservation variety/Landrace | Finland | Norway | + | - |
| **Tvengsberg NordGen** | - | Conservation variety/Landrace | Norway | Norway | - | + |
| **Hankkijan Jussi\*** | - | Population variety | Finland | Russia | + | + |
|  |  | (Selection from 60Co irradiated 'Vjatka) |  |  |  |  |

*\*Rye varieties analysed by both: SSRs and SNPs – Tvengsberg NordGen was the only genotype not included in SSRs analyses; while Mustialan Tvengsberg, Sangaste and Reetta were not included in SNPs analyses. Lo7 and hybrid rye were used as an outgrouping controls.*
*Morphological descriptions of conservation varieties according to UPOV will be available in supplemental data upon publication of a manuscript. The origin of population varieties was found from* Pluto database https://www.upov.int/pluto/en/

Inbred line Lo7 from KWS Lochow GmbH, Germany, and an unnamed hybrid variety from Germany were included as controls in the genetic diversity analyses but were excluded from the AMOVA of the rye population varieties.

In SSR analyses, the number of plants studied from each variety varied between 28-30 (445 individual plants in total). In SNP analyses, the number of plants studied was mostly 6 (in ‘Riihi’ 5) and the total number of individual plants was 59. Varieties ‘Sangaste’, ‘Reetta’ and ‘Mustialan Tvengsberg’ were not included in the SNP analyses nor the controls (SNP data was created in SUSCROP ERANET project, called RYE-SUS). However, Tvengsberg from NordGen (Nordic genebank) was included in the SNP analyses.

### DNA extractions, SSR and SNP genotyping

For SSR analyses, plants were grown in a greenhouse and DNA was extracted from young, fresh rye leaves (at the three-leaf stage) with an E.Z.N.AÓ SP Plant DNA kit (Omega Bio-tek, GA, USA). DNA concentrations were measured using a NanoDrop™ One/OneC Microvolume UV-Vis Spectrophotometer (Thermo Fisher Scientific, MA, USA). For SNP analyses, leaf material was grown at Julius Kühn Institute (JKI), Germany, and sampled according to the instructions of SGS INSTITUT FRESENIUS GmbH, TraitGenetics section (SGS IF TG), Germany, where DNA was extracted.

Eighteen SSRs developed for rye (Hackauf and Wehling 2002; Saal and Wricke 1999) were primarily selected for the diversity study. They were amplified in five different PCR reactions and first tested in a smaller number of samples. Ten SSRs, which were amplified reliably, were polymorphic, and yielded unambiguous allele patterns, were used for the final analyses (Suppl. Table S2). An ABI PRISM® 310 Genetic Analyzer (Applied Biosystems, CA, USA) was used to separate and visualise amplified products. The forward primer of each primer pair was labelled with a fluorescent dye, FAM™ (5-carboxyfluorescein), NED™, VIC™, or PET™. The PCR amplification conditions were as follows: an initial denaturation step of 15 min at 95°C, followed by 35 cycles of 30 s at 94°C, 90 s at the primer specific annealing temperature (Ta, Supplementary Table S2), and 60 s at 72°C, and a final extension step of 30 min at 60°C. Amplifications were performed in a Bio-Rad C1000 Thermal Cycler (Bio-Rad Laboratories, CA, USA).

The PCR amplification was performed in a total volume of 10 μL, containing 5 μL of 2x Qiagen Master Mix from the Qiagen Multiplex PCR Kit (Qiagen, Hilden, Germany), 10 ng genomic DNA, and 2 μM of each primer. PCR products were diluted 1:250 for the ABI runs. Allele size estimation was performed using GeneMapperÓ Software v5.0 (Applied Biosystems, CA, USA). SNP genotyping was performed with the wheat/rye/triticale 25K + 5K Infinium Array (Illumina, CA, USA) at SGS IF TG, Germany. The array was based on 25K wheat array, to which approximately 5K rye markers from the published Rye Infinium array (Haseneyer et al. 2011) were added, together with an additional set of markers from the 600K Affymetrix array (Bauer et al. 2017). A total of 5084 polymorphic SNPs out of 10,168 SNPs were used in the diversity analyses.

### Genetic diversity, clustering and population structure

For the SSR and SNP analyses, the polymorphic information content (PIC) and number of alleles were calculated using CERVUS version 3.0.7 (Kalinowski et al. 2007). The program GenAlEx 6.5 (Peakall and Smouse 2006; Peakall and Smouse 2012) was used to calculate observed heterozygosity (Ho), expected heterozygosity (He), percentage of polymorphic loci, and Shannon’s diversity index (I; Shannon 1948). To further investigate genetic variation between varieties and landraces, analysis of molecular variance (AMOVA) was performed to calculate genetic variance within and among populations. Principal coordinates analysis (PCoA) based on Nei’s genetic distances (Nei 1972) was conducted with GenAlEx (Peakall and Smouse 2012). Statistical significance was tested using 999 permutations. SSR and SNP data were analysed separately, and a dissimilarity matrix based on the Dice coefficient was calculated using DARwin version 6.0.014 (Perrier and Jacquemoud-Collet 2006). An unweighted neighbour-joining (NJ; Saitou and Nei 1987) tree was constructed from the dissimilarity matrix. In addition, an UPGMA tree was generated from the SSR data of rye populations using MEGA software (Kumar et al. 2024).

For Bayesian clustering analyses, STRUCTURE version 2.3.4 (Pritchard et al. 2000) was used. The analysis was run for the most probable number of genetic clusters (K), ranging from 1 to 15, with 10 independent replicates per K. Each run employed a burn-in period of 150,000 iterations followed by 200,000 MCMC (Markov Chain Monte Carlo) repetitions under the admixture model with correlated allele frequencies. The optimal number of clusters (K) was inferred using the ΔK method of Evanno et al. (2005) and determined separately for each dataset, including analyses with and without Lo7, as well as analyses of the hybrid and ‘Tvengsberg’ datasets.

## Results

The 10 selected SSRs for the study amplified a total of 55 alleles (Suppl. Table S2). Their PIC value varied from 0.377 to 0.804 with the mean of 0.56. The number of SNP markers in the study was 5084, and the average PIC value 0.2902 (Suppl. Table S3).

SSRs were 100 % polymorphic in all varieties except in the hybrid variety (90 %) and in Lo7 (inbred line, 0 %, Table 2). The average values without hybrid and Lo7 were: I 1.035, *H*_o_ 0.534, and *H*_e_ 0.583; the total number of alleles varied from 35 (Elvi and Jussi) to 42 (Pääkkölä), the mean being 39. In the SNP analysis the number of polymorphic loci in the 10 varieties studied varied from 83.4 to 90.8 % (Suppl. Table S4), the average values for I, *H*_o_, and *H*_e_ were 0.478, 0.341 and 0.321, respectively. AMOVA revealed that most genetic variation occurred within populations for both marker systems (92%), whereas 8% was attributable to differences among populations. Population differentiation was nevertheless significant for both SSRs (PhiPT = 0.084, P = 0.001) and SNPs (PhiPT = 0.077, P = 0.001). In the dendrogram using SSRs, individual plants (Suppl. Figure S1) did not group together according to varieties but were mainly scattered to several branches. The only exceptions were the identical samples of the inbred line Lo7, and the samples of the hybrid rye, which were mainly grouped in two branches of the tree. In the SNP dendrogram (Supplementary Figure S2), individuals of Tvengsberg (NordGen) and Jussi showed comparatively distinct clustering, while Riihi and Polvijärvi also showed substantial within-accession grouping. Iivo individuals clustered predominantly together, although two individuals occurred in other branches. In contrast, individuals of Joppe, Hermanni and Pääkkölä were interspersed across branches, consistent with substantial shared genetic variation among these accessions. The UPGMA tree of SSR results (Figure 1) clearly indicated that Tvengsberg (NordGen), the hybrid variety, and Lo7 formed separate branches, whereas the remaining varieties clustered into a larger group.

**Table 2.** Diversity indices of SSRs in 15 rye landraces/varieties.

| **Population** | **Number of individuals (N)** | **Number of alleles (n)** | **No. of private alleles** | **Shannon's Information Index (I)** | **Observed heterozygosity (Ho)** | **Expected heterozygosity (He)** | **Polymorphic loci (%)** | **Inbreeding coefficient (FIS)** | **Hardy-Weinberg exact test** |
|---|---|---|---|---|---|---|---|---|---|
| **Aunus** | 30 | 41 | 1 | 1.071 | 0.557 | 0.588 | 100 | 0.0078 | ** |
| **Elvi** | 30 | 35 | 0 | 1.025 | 0.568 | 0.586 | 100 | 0.0477 | * |
| **Hermanni** | 28 | 40 | 0 | 1.080 | 0.500 | 0.601 | 100 | 0.1909 | *** |
| **Hybrid** | 30 | 25 | 0 | 0.694 | 0.567 | 0.433 | 90 | -0.3078 | *** |
| **Iivo** | 30 | 40 | 2 | 1.029 | 0.507 | 0.570 | 100 | 0.1316 | *** |
| **Joppe** | 29 | 42 | 2 | 1.074 | 0.529 | 0.588 | 100 | 0.1135 | ** |
| **Jussi** | 30 | 35 | 0 | 0.924 | 0.443 | 0.535 | 100 | 0.2023 | *** |
| **Lo7** | 30 | 10 | 1 | 0.000 | 0.000 | 0.000 | 0 | NA<sup>a</sup> |  |
| **Polvijärvi** | 30 | 39 | 0 | 1.047 | 0.582 | 0.599 | 100 | 0.0600 | *** |
| **Pääkkölä** | 30 | 42 | 0 | 1.056 | 0.564 | 0.590 | 100 | 0.0656 | *** |
| **Reetta** | 30 | 39 | 1 | 1.042 | 0.534 | 0.596 | 100 | 0.1160 | *** |
| **Riihi** | 29 | 37 | 1 | 1.026 | 0.564 | 0.585 | 100 | 0.0620 | ** |
| **Sangaste** | 30 | 37 | 1 | 0.986 | 0.485 | 0.566 | 100 | 0.1570 | *** |
| **Taavetti** | 29 | 41 | 0 | 1.085 | 0.594 | 0.604 | 100 | 0.0300 | *** |
| **Mustialan Tvengsberg** | 30 | 36 | 1 | 1.006 | 0.519 | 0.567 | 100 | 0.1130 | ** |
|  | **Total**<br>445 | **Average**<br>36 | **Total**<br>10 | **Average**<br>0.943 | **Average**<br>0.501 | **Average**<br>0.534 | - | - | - |
| **Average excluding Hybrid and Lo7** |  |  |  | 1.035<sup>b</sup> | 0.534<sup>b</sup> | 0.583<sup>b</sup> |  |  |  |

*Probability of the multilocus Hardy-Weinberg exact test. \*P < 0.05, \*\*P < 0.01, \*\*\*P < 0.001. ^a All individuals are genetically identical. ^b Excluding the hybrid cultivar and Lo7.*
*NA, not applicable because all individuals were genetically identical.*

**Table 3.** Analysis of molecular variance (AMOVA) among and within rye populations: (a) 13 population varieties genotyped with 10 SSR markers, excluding the inbred line Lo7 and hybrid control; and (b) 10 accessions genotyped with 5084 SNP markers.

| a) |  |  |  |  |  |
|---|---|---|---|---|---|
| **Source** | **df** | **SS** | **MS** | **Variance components** | **Total (%)** |
| **Among Populations** | 12 | 299.767 | 24.981 | 0.617 | 8% |
| **Within Populations** | 372 | 2490.635 | 6.695 | 6.695 | 92% |
| **Total** | 384 | 2790.403 |  | 7.313 | 100% |
| **Stat** | Value | P(rand >= data) |  |  |  |
| **PhiPT** | 0.084 | 0.001 |  |  |  |
| b) |  |  |  |  |  |
| **Source** | **df** | **SS** | **MS** | **Variance components** | **Total (%)** |
| **Among Populations** | 9 | 26355.889 | 2928.432 | 163.346 | 8% |
| **Within Populations** | 49 | 96283.467 | 1964.969 | 1964.969 | 92% |
| **Total** | 58 | 122639.356 |  | 2128.315 | 100% |
| **Stat** | Value | P(rand >= data) |  |  |  |
| **PhiPT** | 0.077 | 0.001 |  |  |  |

*Probability, P (rand >= data), for PhiPT is based on 999 permutations across the full data set.; PhiPT = AP / (WP + AP) = AP / TOT.; Key: AP = estimated variance among populations, WP = estimated variance within populations.*

**Figure 1.**
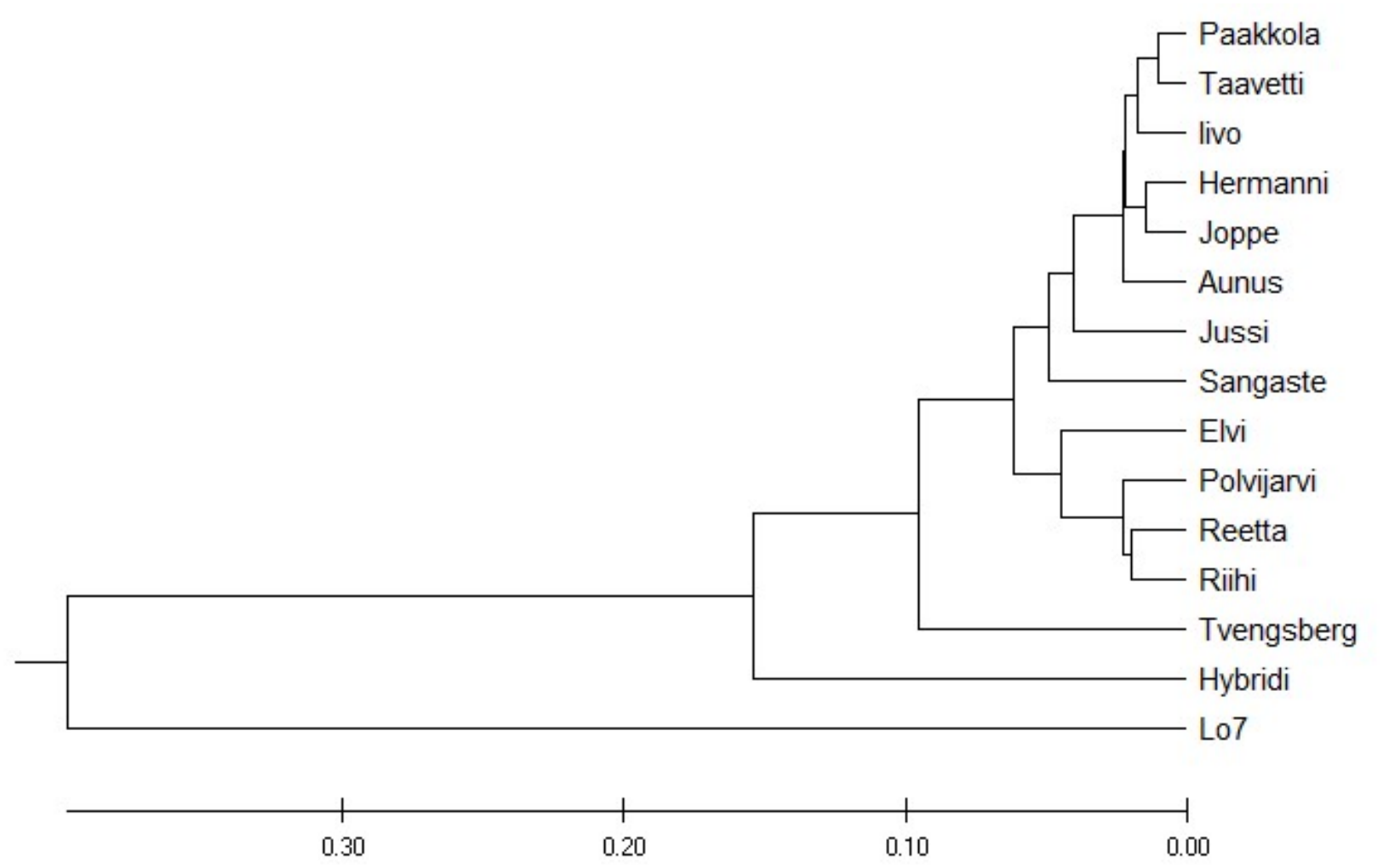
UPGMA clustering of Nordic rye germplasm based on SSR markers separates Lo7, hybrid rye and Tvengsberg from the main Nordic rye gene pool and identifies a closely related Reetta-Riihi-Polvijärvi-Elvi group.

Genetic distances between populations were visualised with PCoA (Figures 2A and 2B). Similar patterns were observed in the SSR-based UPGMA dendrogram (Figure 1), in which Lo7 and the hybrid were strongly differentiated from the population varieties, while Tvengsberg was also differentiated from the remaining Nordic rye material. Within this cluster, Elvi, Polvijärvi, Reetta, and Riihi, grouped together and formed their own subcluster. When the controls were excluded (Figures 2A, 2B), the first two axes explained 74.8% of (SSR) or 11.68% (SNP) of the total variation, respectively. In the first SSR-based PCoA Figure 2A, 13 populations were used as samples, and in the second SNP-based PCoA Figure 2B, 59 samples were analysed. ‘Mustialan Tvengsberg’ was the most distinct of all the other varieties.

**Figure 2.**
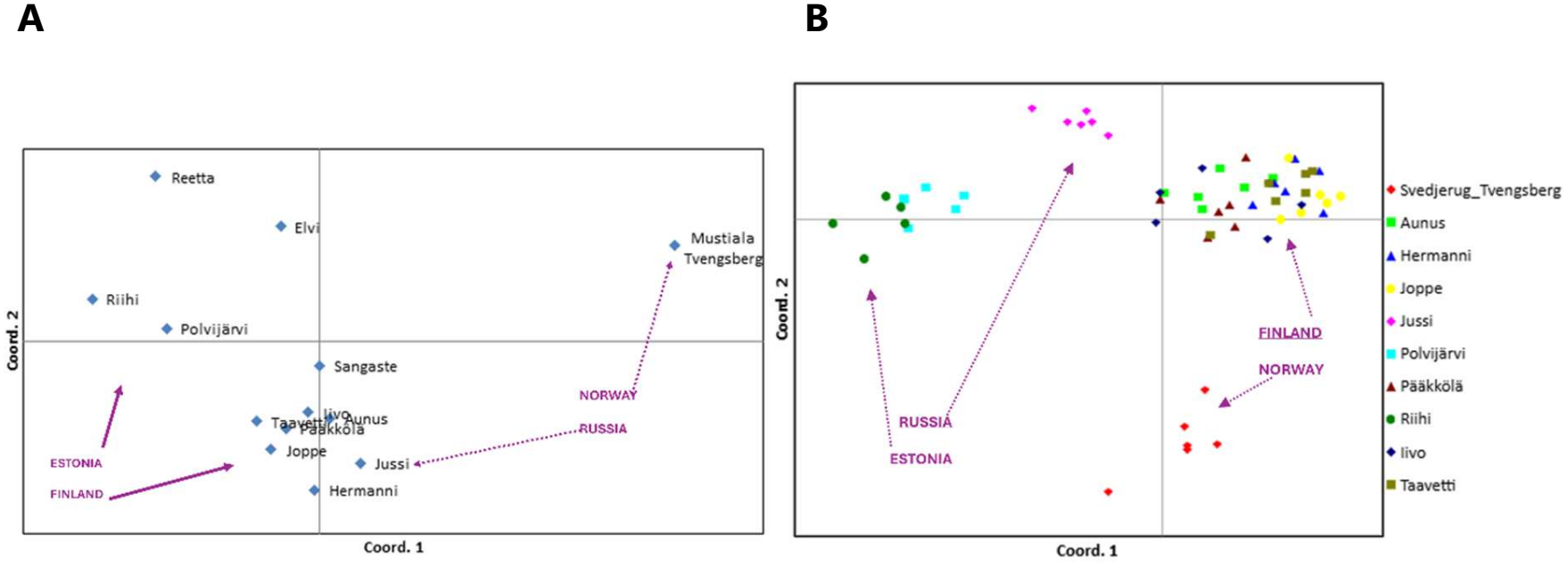
Principal coordinate analyses reveal geographically associated differentiation within the Nordic rye germplasm (Finnish, Estonian, Norwegian, and Russian) within a largely shared Nordic rye gene pool. **(A)** SSR-based PCoA of 13 rye population varieties using approximately 30 plants per variety. Each point represents the genetic centroid of a population. Tvengsberg is clearly separated from the remaining accessions, while Elvi, Reetta, Riihi and Polvijärvi form a distinct cluster. Aunus, Hermanni, Iivo, Joppe, Pääkkölä and Taavetti group closely together, with Sangaste and Jussi occupying intermediate positions. **(B)** SNP-based PCoA based on 5084 markers genotyped in individual plants from ten rye accessions. Tvengsberg forms a highly differentiated cluster, whereas Jussi occupies a distinct position along the second principal coordinate. Riihi and Polvijärvi cluster closely together, while Aunus, Hermanni, Iivo, Joppe, Pääkkölä and Taavetti show substantial overlap. Both marker systems identified Tvengsberg as the most differentiated Nordic accession. In the SSR analysis, Elvi, Reetta, Riihi and Polvijärvi also showed close genetic relationships, whereas the SNP analysis supported close genetic similarity between Riihi and Polvijärvi. The extensive overlap among most remaining accessions is consistent with AMOVA results showing that 92% of the genetic variation resides within populations and only 8% among populations, indicating extensive shared ancestry and admixture within Nordic-Baltic rye germplasm.

Clustering of rye populations with SSR results was analysed using STRUCTURE. In the first analysis (Figure 3), all samples were included (n = 445; approximately 30 seeds per population variety). The strongest statistical support was obtained for K = 2 (ΔK = 242). The second-highest support was observed for K = 5, although with a substantially lower ΔK value (ΔK = 8.9845). At K = 2, Lo7 and the hybrid formed one genetic cluster, whereas all other varieties were assigned predominantly to a second cluster. Thus, Lo7 and the hybrid were clearly differentiated from the remaining rye accessions, which formed a common genetic group.

**Figure 3.**
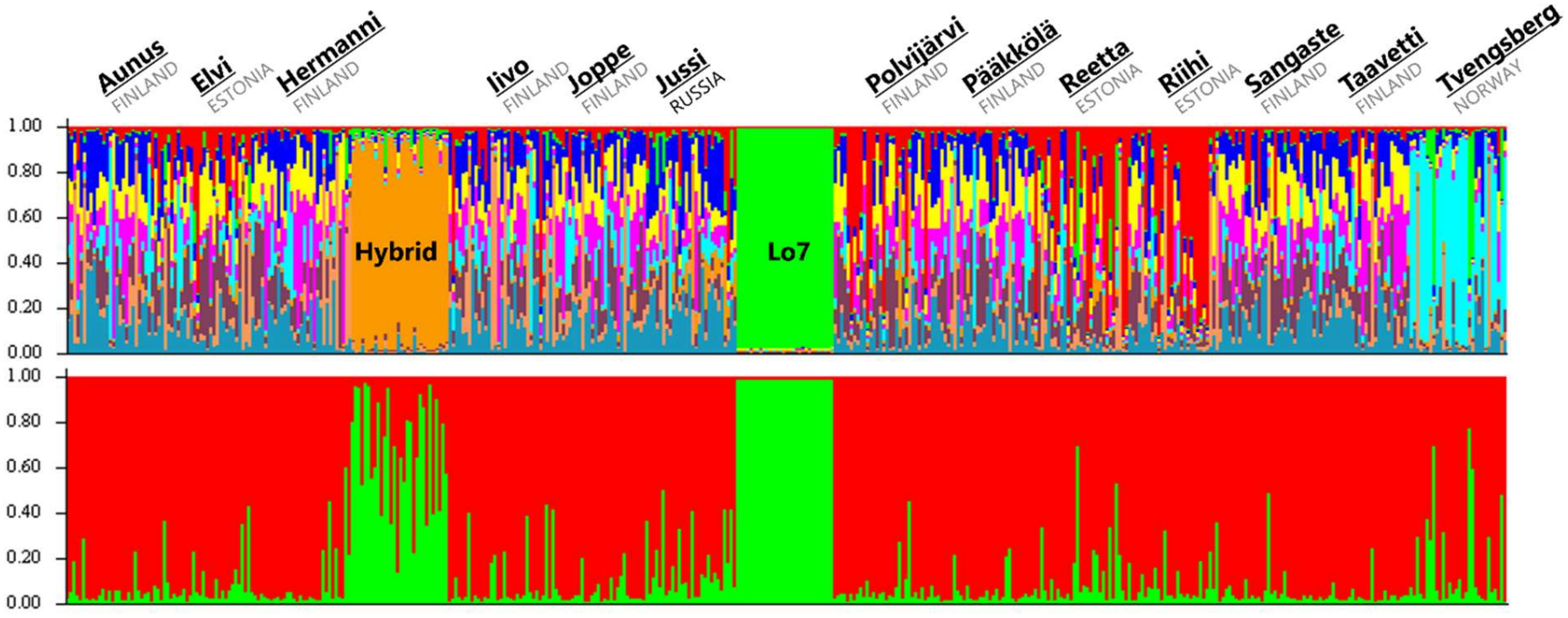
STRUCTURE analysis of the complete rye dataset (n = 445) identifies two major ancestry groups (K = 2; ΔK = 242), with Hybrid and Lo7 assigned predominantly to one cluster and all remaining accessions to the other. Each vertical bar represents an individual plant (30 plants per accession/population cultivar). Colors indicate estimated ancestry coefficients. Accession names are shown in bold above each group of individuals, and the putative country of origin is indicated above the accession name. Hybrid and Lo7 are labelled directly within the STRUCTURE plot because they form distinct ancestry groups at K = 2. The K = 2 solution received the strongest support (ΔK = 242), whereas all alternative K values showed substantially lower support (ΔK < 9).

To investigate genetic differentiation within the Nordic rye material, Lo7 and the hybrid were excluded from the subsequent analysis (Figure 4). The second STRUCTURE analysis included only the Nordic rye accessions (n = 385; approximately 30 seeds per population variety). Both K = 2 and K = 3 received the strongest statistical support. At K = 2 (ΔK = 29), ‘Tvengsberg’ formed a distinct genetic cluster, while all other varieties were assigned mainly to a second cluster (results not shown).

**Figure 4.**
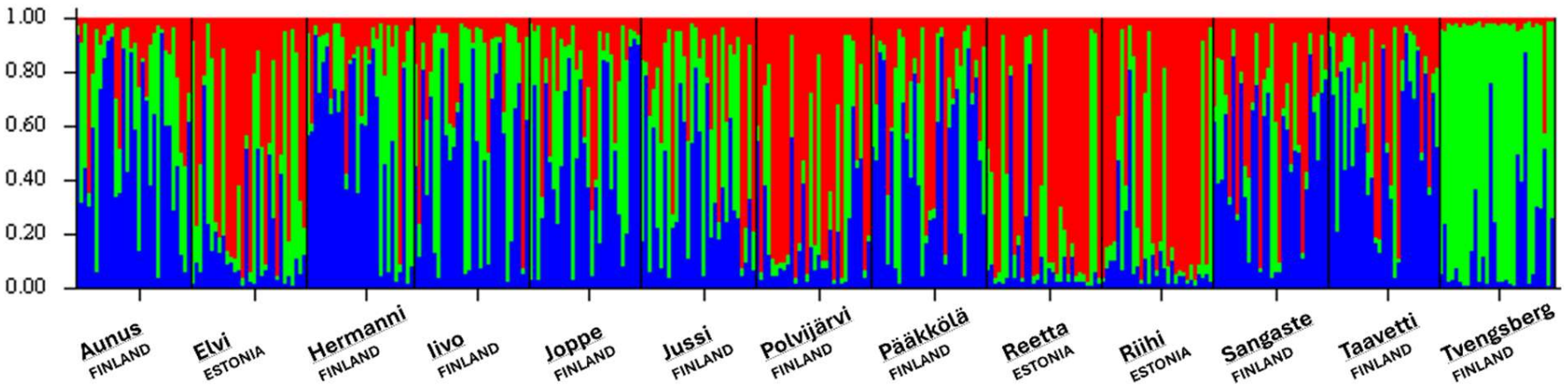
STRUCTURE analysis of Nordic rye accessions excluding Hybrid and Lo7 (n = 385; 13 population varieties; appr. 30 plants per variety) at K = 3. Each bar represents an individual plant, with colors indicating ancestry proportions. Variety names are shown in bold and putative countries of origin are indicated above the corresponding groups. The K = 2 (ΔK = 29) and K = 3 (ΔK = 27) models received the strongest support. At K = 2, Tvengsberg formed a distinct cluster, while all remaining accessions grouped together. At K = 3, three major ancestry groups were resolved: (i) Tvengsberg, (ii) Reetta-Riihi-Polvijärvi (with partial contribution from Elvi), and (iii) Aunus-Hermanni-Iivo-Joppe-Pääkkölä-Taavetti-Sangaste. Extensive admixture among varieties suggests shared ancestry. Because Tvengsberg remained highly distinct in both solutions, it was excluded from subsequent analyses to reveal finer-scale structure among the remaining Nordic rye accessions.

At K = 3 (ΔK = 27), three genetic groups could be distinguished. The first group consisted primarily of Tvengsberg individuals. The second group included ‘Reetta’, ‘Polvijärvi’ and ‘Riihi’ with some contribution from ‘Elvi’. The third group comprised ‘Aunus’, ‘Hermanni’, ‘Iivo’, ‘Joppe’, ‘Pääkkölä’, and ‘Taavetti’ with contribution from ‘Sangaste’. These results indicate a moderate level of genetic structuring within the Nordic rye material.

To further examine population structure among the Nordic rye accessions, ‘Tvengsberg’ was excluded from an additional STRUCTURE analysis (Figure 5).

**Figure 5.**
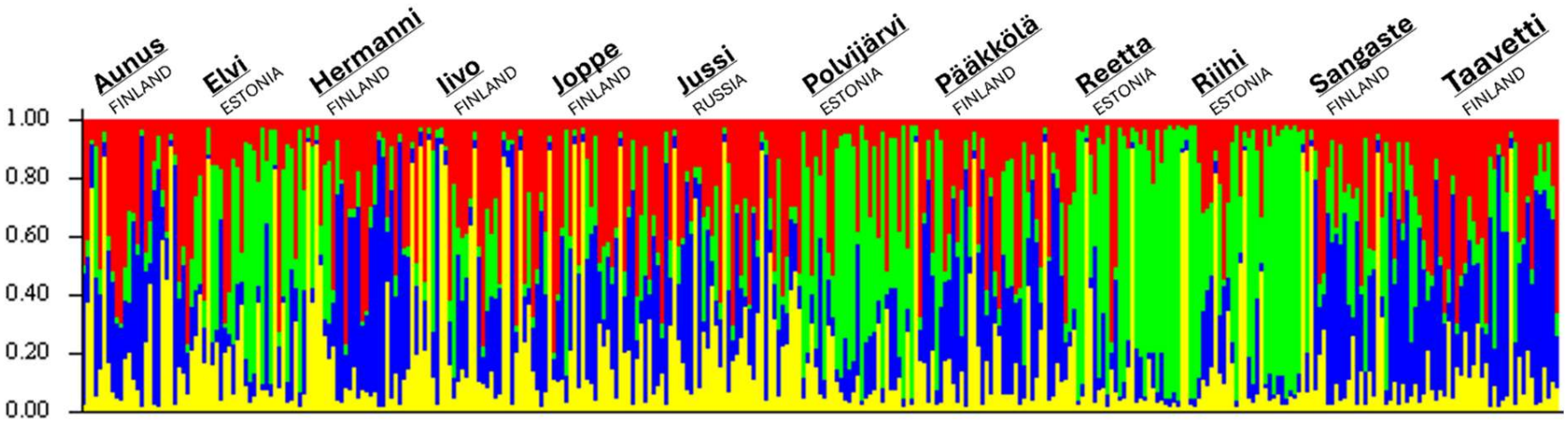
Following exclusion of Mustialan Tvengsberg, STRUCTURE analysis of the remaining Nordic rye accessions (n = 355; approximately 30 plants per accession) identified four ancestry components (the strongest support was obtained for K = 4; ΔK = 7.8). Each vertical bar represents an individual plant and colors indicate ancestry proportions. Accession names are shown in bold above each group. Elvi, Reetta and Riihi showed high membership in a common ancestry component, with Polvijärvi displaying substantial assignment to the same cluster. Hermanni, Sangaste and Taavetti were predominantly associated with a second ancestry component, whereas Iivo showed elevated assignment to a third component. In contrast, Jussi exhibited relatively balanced contributions from multiple ancestry components, suggesting a more admixed genetic background.

Following exclusion of Tvengsberg, several ancestry patterns became apparent among the remaining rye populations. Elvi, Reetta and Riihi shared a prominent ancestry component, also present in Polvijärvi, whereas Hermanni, Sangaste and Taavetti were characterized by another component that was also detected in parts of Aunus and Joppe. Other ancestry components were more broadly distributed across the material, including one strongly represented in Iivo and a background component occurring in most populations. These patterns suggest differentiation among particular groups of populations against a background of extensive shared ancestry.

Jussi was notable for its heterogeneous ancestry composition, with contributions from several of the ancestry components observed elsewhere in the material rather than assignment predominantly to a single cluster. This pattern is consistent with a comparatively admixed genetic background. More broadly, the STRUCTURE results agree with the AMOVA in indicating that detectable population structure occurs within a largely shared gene pool containing substantial within-population diversity.

## Discussion

Studies based on isoenzyme, phenotypic and molecular markers have consistently shown that in cross-pollinating species genetic variation is predominantly distributed within rather than among populations. Similar results have been reported for Finnish landraces, varieties and gene bank accessions, where up to 86% of the diversity was found within accessions (Persson and Bothmer 2002; Persson et al. 2006; Targońska et al. 2016; Hagenblad et al. 2016). Consistent with these findings, our AMOVA showed that 92% of the total genetic variation was attributable to differences within populations, whereas only 8% was explained by variation among populations.

In Persson and Bothmer (2002) and Persson et al. (2006), Swedish and Finnish landraces showed higher genetic variability than landraces from Germany, Poland and other Nordic countries, although this may partly reflect differences in sample size. Finnish, Swedish and Norwegian accessions were genetically more similar to each other than to accessions from Germany and Poland. The close genetic relationship between Finnish and Swedish landraces likely reflects their shared history, as Finnish rye landraces were introduced to central Sweden by Finnish settlers during the sixteenth and seventeenth centuries (Hagenblad et al. 2016). Genetic similarity among Finnish landraces may also result from a common ancestry and historical gene flow through hybridization between populations (Persson and Bothmer 2002). In contrast, Ma et al. (2004), using RAPD markers, found Finnish varieties to be genetically closer to Central and Northern European varieties, particularly those from Germany and Poland, suggesting a shared genetic background and adaptation to similar environmental conditions. Their results also supported the Russian genetic origin of the rye cultivar ‘Hankkijan Jussi’.

Recent molecular studies further support the existence of a broadly shared Fennoscandian rye gene pool. Using 22 SSR markers, Targońska et al. (2016) showed that rye landraces often had mixed ancestry across wide geographical areas and that breeding lines and modern varieties possessed a narrower genetic base than gene bank accessions, a pattern also reported by Chikmawati et al. (2012) and Bolibok-Brągoszewska et al. (2014). Hagenblad et al. (2016) analysed approximately 570 SNP markers in nearly 450 rye accessions, found that accessions from northern Fennoscandia clustered together with other Fennoscandian and some Eastern European genotypes, whereas Russian landraces formed a more distinct group. Similarly, Larsson et al. (2019) suggested that Fennoscandian rye landraces constitute a single major metapopulation that has remained largely stable for at least 350 years. Historical records indicate that rye seed was exported from Finland to the Baltic region and Russia during medieval times (Ahokas 2009), which may also have contributed to the genetic similarity among northern and eastern landrace populations.

Our results support a broadly shared Fennoscandian rye gene pool while revealing reproducible regional differentiation. Both SSR and SNP analyses revealed geographically associated genetic differentiation, although substantial overlap and admixture occurred among accessions. The Finnish landraces ‘Aunus’, ‘Hermanni’, ‘Iivo’, ‘Joppe’, ‘Pääkkölä’, and ‘Taavetti’ generally showed similar genetic profiles, particularly in the SNP analysis. Admixture among accessions indicates shared ancestry and historical gene flow, whereas the consistent regional structure shows that geographic differentiation has persisted despite rye’s outcrossing breeding system and historical seed exchange. This interpretation is also consistent with the AMOVA results, which showed that most variation occurred within populations.

The SSR clustering further identified relationships consistent with accession histories. The genetic proximity observed among Elvi, Reetta, Riihi and Polvijärvi suggests shared ancestry or historical gene flow, although the available pedigree information does not presently allow the origin of this relationship to be resolved. In contrast, the separate position of ‘Hankkijan Jussi’ agrees with its documented origin as a selection from the 60Co-irradiated Russian variety ‘Vjatka’ (Ma et al. 2004).

‘Tvengsberg’ landrace was the clearest exception to the general overlap among Fennoscandian accessions. The independently sampled Mustialan and NordGen accessions each occupied a distinct position in the SSR and SNP datasets, respectively, providing convergent evidence for differentiation of Tvengsberg from the remaining material. This pattern may reflect ‘Tvengsberg’’s history: the landrace was recovered from an old drying barn in Finnskogen, a forest region on the Norwegian-Swedish border, reintroduced as a separate population, and remained geographically isolated from Finnish rye populations for centuries (Svedjerug: Natural History Museum and Botanical Garden 2026). Genetic drift, local adaptation, and the recently very small population size may therefore have contributed to its divergence. Nevertheless, its historical origin suggests that ‘Tvengsberg’ represents a differentiated branch of the traditional Finnish slash-and-burn rye gene pool rather than an independent origin. The inbred line Lo7 and the hybrid rye were also clearly distinct and more genetically uniform than the open-pollinated populations, supporting their use as outgroup controls.

Sequential STRUCTURE analyses revealed hierarchical genetic structure within the rye collection. When all accessions were included, the major partition separated Hybrid and Lo7 from the remaining materials. Following removal of these highly differentiated accessions, Tvengsberg emerged as the primary source of genetic differentiation among Nordic ryes. Excluding Tvengsberg further exposed finer-scale structure within the Nordic germplasm, revealing groups represented by Reetta-Riihi-Polvijärvi-Elvi, Hermanni-Sangaste-Taavetti, and Iivo-related accessions. These results suggest that genetic diversity within Nordic rye is organized hierarchically, with successively finer ancestry patterns becoming apparent as dominant sources of variation are removed.

Marker informativeness differed substantially between the datasets. The 10 SSR markers detected 55 alleles and had PIC values of 0.377–0.804 (mean 0.56), whereas the mean PIC of the 5084 SNP markers was 0.290. The high informativeness of rye SSR markers agrees with Targońska et al. (2016) and was also evident in our STRUCTURE analyses.

As multi-allelic markers, SSRs captured allelic richness and rare variants more effectively than bi-allelic SNPs and were therefore particularly informative for diversity within heterogeneous landrace populations. SNPs nevertheless provide efficient genome-wide and high-throughput characterization, although panels developed mainly from elite breeding material may miss some landrace variation.

## Conclusions

The high genetic diversity within Finnish rye landraces, together with the regional differentiation detected among Nordic accessions, highlights the value of these populations for conservation and future breeding. The concordant patterns revealed by SSR and SNP markers show that a broadly shared Fennoscandian gene pool can retain geographically structured variation despite historical seed exchange and rye’s outcrossing breeding system. As climate change increases environmental variability and the frequency of abiotic and biotic stresses, these genetically diverse landrace populations may provide useful sources of adaptation. Their conservation is therefore important not only for preserving cultural heritage but also for securing genetic resources for sustainable rye production and breeding.

## Acknowledgements

We thank Marja-Riitta Arajärvi, Sirpa Moisander and Anneli Virta (Luke) for technical assistance. We thank Boreal Plant Breeding Ltd., KWS SAAT SE & Co. KGaA, Jorma Aaltonen, Antti Laine, Matti Pappinen, Sakari Raiskio, Hannu Snellman, Lauri Takala, Häme University of Applied Sciences and Metsähallitus for providing rye material, and Minna-Maarit Kytöviita and Annika Michelson for coordinating its collection and delivery.

This research was supported by the EU Horizon 2020 ERA-NET Cofund SusCrop RYE-SUS project (Grant No. 771134) and the Finnish Ministry of Agriculture and Forestry through the *Diversity to* Fields (MoPe) project.

AI tools were used for language editing and manuscript structure.

## References

Ahokas, H. 2009. The exported winter rye form evolved in co-cultivation with spring-sown seasonal crops in Finland where the slash-and-burn and riihi traditions selected against the rachis brittleness of the weedy rye. Interdisciplinary Biology, Agriculture, Linguistics and Agriculture 4: 1–153.

Ahokas, H. 2012. Crop evolution under fire: the past cultivation with sequential kytö burning selected against the shattering weedy forms and comparison between Finnish kytö and Ethiopian guie. Helsinki: Kave. 194 p. (Interdisciplinary Biology, Agriculture, Linguistics and Antiquities, No. 5). ISBN 978-951-98445-6-5.

Bolibok-Brągoszewska, H., Targońska, M., Bolibok, L., Kilian, A., & Rakoczy-Trojanowska, M. 2014. Genome-wide characterization of genetic diversity and population structure in Secale. BMC Plant Biology 14: 184.

Chikmawati, T., Miftahudin, M., Skovmand, B., & Gustafson, J.P. 2012. Amplified fragment length polymorphism-based genetic diversity among cultivated and weedy rye (Secale cereale L.) accessions. Genetic Resources and Crop Evolution 59: 1743–1752.

Evanno, G., Regnaut, S. & Goudet, J. (2005). Detecting the number of clusters of individuals using the software STRUCTURE: a simulation study. Molecular Ecology 14: 2611–2620.

Evans, G.M. 1976. Rye, Secale cereale (Gramineae-Triticinae). In: Simmonds, NW (ed.), Evolution of crop plants: 108-111. Longman Inc., New York, the United States of America.

Hackauf, B. & Wehling, P. 2002. Identification of microsatellite polymorphisms in an expressed portion of the rye genome. Plant Breeding 121: 17–25.

Hagenblad, J., Hugo, O., Forsberg, N. & Leino, M. 2016. Geographical distribution of genetic diversity in Secale landrace and wild accessions. BMC Plant Biology 16:23

Haseneyer, G., Schmutzer, T. Seidel, M., Zhou, R., Mascher, M., Schön, C.C., Taudien, S., Scholz, U., Stein, N., Mayer, K.F.X. & Bauer, E. 2011. From RNA-seq to large-scale genotyping - genomics resources for rye (Secale cereale L.). BMC Plant Biology 11: 131.

Kalinowski, S.T., Taper, M.L., & Marshall, T.C. 2007. Revising how the computer program CERVUS accommodates genotyping error increases success in paternity assignment. Molecular Ecology 16: 1099–1106.

Kumar, S., Stecher, G., Suleski, M., Sanderford, M., Sharma, S., and Tamura, K. (2024). Molecular Evolutionary Genetics Analysis version 12 for adaptive and green computing. Molecular Biology and Evolution 41: 1–9.

Larsson, P., Oliveira, H.R., Lundström, M., Hagenblad, J., Lagerås, P. & Leino, M.W. 2019. Population genetic structure in Fennoscandian landrace rye (Secale cereale L.) spanning 350 years. Genetic Resources and Crop Evolution 66: 1059–1071.

Ma, R., Yli-Mattila, T. & Pulli, S. 2004. Phylogenetic relationships among genotypes of worldwide collection of spring and winter ryes (Secale cereale L.) determined by RAPD-PCR markers. Hereditas 140: 210–221.

Nei, M. 1972. Genetic distance between populations. The American Naturalist 106: 283–292.

Peakall, R., & Smouse, P.E. 2006. GENALEX 6: genetic analysis in Excel. Population genetic software for teaching and research. Molecular Ecology Notes 6: 288–295.

Peakall, R. & Smouse, P.E. 2012. GenAlEx 6.5: genetic analysis in Excel. Population genetic software for teaching and research – an update. Bioinformatics 28: 2537–2539.

Perrier, X. & Jacquemoud-Collet, J.P. 2006. DARwin software. CIRAD, Montpellier, France.

Persson, K. & Bothmer, R. von, 2002. Genetic diversity amongst landraces of rye (Secale cereale L.) from northern Europe. Hereditas 136: 29–38.

Persson, K., Bothmer, R. von, Gullord, M. & Gunnarsson, E. 2006. Phenotypic variation and relationships in landraces and improved varieties of rye (Secale cereale L.) from northern Europe. Genetic Resources and Crop Evolution 53: 857–866.

Pritchard, J.K., Stephens, M. & Donnelly, P. 2000. Inference of population structure using multilocus genotype data. Genetics 155: 945–959.

Saal, B. & Wricke, G. 1999. Development of simple sequence repeat markers in rye (Secale cereale L.). Genome 42: 964–972.

Saitou, N. & Nei, M. 1987. The neighbor-joining method: a new method for reconstructing phylogenetic trees. Molecular Biology and Evolution 4: 406–425.

Schreiber, M., Himmelbach, A., Börner, A., and Mascher, M. 2018. Genetic diversity and relationship between domesticated rye and its wild relatives as revealed through genotyping-by-sequencing. Evolutionary Applications 12: 66–77.

Shannon, C.E. 1948. A mathematical theory of communication. Bell System Technical Journal 27: 379–423, 623–656.

Targońska, M., Bolibok-Brągoszewska, H. & Rakoczy-Trojanowska, M. Assessment of Genetic Diversity in Secale cereale Based on SSR Markers. Plant Mol Biol Rep 34, 37–51 (2016). 10.1007/s11105-015-0896-4

